# Direct quantification of the metabolic heat output of individual *Drosophila* brains

**DOI:** 10.1101/2025.08.08.669302

**Authors:** Kanishka Panda, Rohith Mittapally, Qianqian Chen, Akshay Bhaskaran, Pramod Reddy, Edgar Meyhofer, Swathi Yadlapalli

## Abstract

Quantitative insights into brain metabolism are essential for advancing our understanding of energy dynamics in the brain. However, current approaches for tracking brain metabolism, metabolic profiling and respirometry, provide only static snapshots of metabolite levels or lack the required resolution. Here, we develop a novel nanowatt-resolution biocalorimeter capable of real-time continuous measurements of heat output to quantitatively measure the metabolism of individual live *Drosophila melanogaster* brains and investigate how sex, genotype, age, and disease affect brain metabolism. We show for the first time that female brains, across multiple wild-type genotypes, exhibit a significantly higher metabolic rate (∼10%) than male brains at a young age (<10 days old) and follow distinct metabolic trajectories across the lifespan. We also find that *parkin* mutants, a genetic model for Parkinson’s disease, exhibit a ∼15% reduction in brain metabolic output relative to controls, revealing that defective mitophagy due to *parkin* deficiency affects brain metabolism. Furthermore, we measure the metabolic rate of reproductive tissues of *Drosophila*, highlighting the broad applicability of our biocalorimeter. Together, these advances open new avenues for investigating how tissue-specific metabolism is impacted by aging, neurodegeneration, and disease states.

**Teaser:** Direct measurement of metabolic rate of individual *Drosophila* brains to investigate how sex, genotype, age, and disease affect brain metabolism.

## INTRODUCTION

The adult human brain, despite comprising only 2% of body weight, consumes 20% of resting energy to drive complex behaviors like decision-making (*1*). This exceptionally high energy demand renders the brain particularly vulnerable to disruptions in energy production and flow. Dysregulated brain metabolism has been implicated in numerous neurodegenerative diseases, including Alzheimer’s (*2*), Parkinson’s (*3*), as well as age-related cognitive decline (*4, 5*). Understanding the energy dynamics in the brain is critical for developing therapeutic interventions. However, current tools for studying cellular bioenergetics in live, intact brains—particularly in small model organisms—remain very limited.

Current approaches for investigating brain metabolism, such as metabolic profiling (*6-11*), and respirometry (*12, 13*), each have critical drawbacks. For example, techniques like metabolic profiling provide only static snapshots of metabolite levels, fail to capture real-time energy fluxes, and typically require destructive sample preparation, precluding long-term metabolic measurements. Respirometry, while being effective in assessing whole-organism oxygen consumption (*12, 13*) or large cell populations or isolated mitochondria (*14*), lacks the sensitivity required for individual tissues from model organisms. As a result, there is a critical need for developing new tools capable of non-invasive quantification of real-time metabolic activity in live brain tissue under physiological and pathophysiological conditions.

Direct calorimetry overcomes the above challenges by measuring metabolic heat output (*15, 16*)—a universal byproduct of cellular energy transduction (*17, 18*)— providing a label-free, integrative view of total energy flux. Recent advances in biocalorimetry have made it possible to study dynamic metabolic processes during early embryonic development (*19-22*), detect age-related metabolic decline in whole *model organisms such as Drosophila melanogaster* (*23*) and *C. elegans* (*24*), and measure heat output from single *Tetrahymena thermophila* cells (*25*). Despite these technical breakthroughs, implementing direct calorimetry to individual, live tissues such as the brain remains challenging, as it requires both sustained viability of tissues or brains and high sensitivity for detecting dynamic changes in metabolism.

Brain explant cultures offer a powerful model system for studying neural activity, development, and disease in a controlled yet physiologically relevant context (*26, 27*). In these preparations, neural tissues are carefully dissected and maintained *ex vivo* in specialized media that preserve their complex cellular architecture and organotypic properties. Brain explants are widely used to study a wide variety of biological phenomena—for instance, rodent hippocampal slices to investigate learning and memory (*26*), and adult retinal explants for neural development and axon guidance (*27*). These explant models provide a critical bridge between *in vivo* experiments and dissociated cell cultures, which makes them ideally suited for probing tissue-level bioenergetics, but their small size and fragility demand highly sensitive, non-invasive tools for real-time metabolic assessment. To date, direct measurements of metabolic heat in live brain explants have not been attempted.

Here, we have developed a high-resolution biocalorimeter capable of measurements with nanowatt sensitivity and a ∼40 s response time while preserving tissue viability via continuous buffer perfusion (∼4 μl/min) that enables, for the first time, direct, real-time measurements of metabolic output from individual, live *Drosophila melanogaster* brains. We validated this technique by both measuring the metabolic output of individual brains and by demonstrating how metabolic activity is affected upon exposure to mitochondrial inhibitors, which suppressed brain metabolism by 25–80%. Further, we uncovered sex-, genotype-, age-, and disease-dependent differences in brain metabolism. Specifically, we discovered that young female brains showed significantly higher metabolic output than males, a difference that diminished with increasing age. In *parkin* mutants modeling Parkinson’s disease, homozygous mutant brains exhibited ∼15% reduced metabolic output relative to heterozygotes. Extending the method to reproductive tissues, we found the metabolic output from individual ovaries and testes to be ∼2.5-fold lower than that of the brain, highlighting the brain’s high energy demands. This approach provides a powerful new tool for real-time metabolic assessment in small, intact brains of model organisms, enabling novel investigations and mechanistic insights into brain energetics, aging, and disease. Beyond brains, this method can be extended to tissue organoids and drug screening platforms, broadening its impact on studies of metabolism, therapeutic development, and age-related disorders.

## RESULTS

### Description of nanowatt resolution biocalorimeter

We developed a highly sensitive nanowatt resolution calorimeter capable of making metabolic measurements from individual *Drosophila* brains. The calorimeter comprises of a vacuum-suspended, capillary-based sensor, i.e. sensing capillary that is housed within two nested shields (Fig. 1A and fig. S1A), which along with μTorr levels of vacuum, μK resolution temperature sensing and PID-based active temperature control, help achieve nanowatt resolution calorimetry. The sensing capillary (Fig. 1B) consists of a cylindrical tube made of Teflon Perfluoroalkoxy resin (PFA), used for its inertness and low thermal conductivity (*28*). The capillary’s dimensions (750 μm inner and 1 mm outer diameter) are selected to accommodate brains and other tissues of *Drosophila* fruit flies, with sizes up to ∼750 μm (*29*) along their largest dimension. A precision thermistor mounted at the center along the length of the capillary acts as a thermometric sensor for detecting temperature changes caused by metabolic heat outputs from *Drosophila* brains. The sensing capillary is suspended across the open side of the inner shield (IS) (see Methods). The anchor points of the capillary (i.e., ends of the suspended part of the capillary tube) are in excellent thermal contact with the IS, which is stabilized to be within ±1 μK of the set point (∼295 K) over 10 hours, as shown by us before (*30*). The sensing capillary further extends out of the two shields and is connected to a syringe pump and a reservoir (Fig. 1A), necessary for loading and unloading brain samples and maintaining continuous flow of a buffer solution during metabolic measurements. A stopper is inserted from one end of the capillary to locate the brain near the thermistor (Fig. 1B). The stopper is made of an optical glass fiber that also acts as a light source for imaging the sample, together with optical windows assembled on the outer shield (OS) and an inverted optical microscope (Zeiss Axiovert 200 with a 10X objective). An identical matching capillary is mounted close to the sensing capillary to enable common-mode cancellation of background thermal drifts, helping enhance the heat power resolution by improving the stability of the temperature signal (*30*). The design and fabrication of the calorimeter is described further in Methods and in fig. S1.

**Fig. 1:**
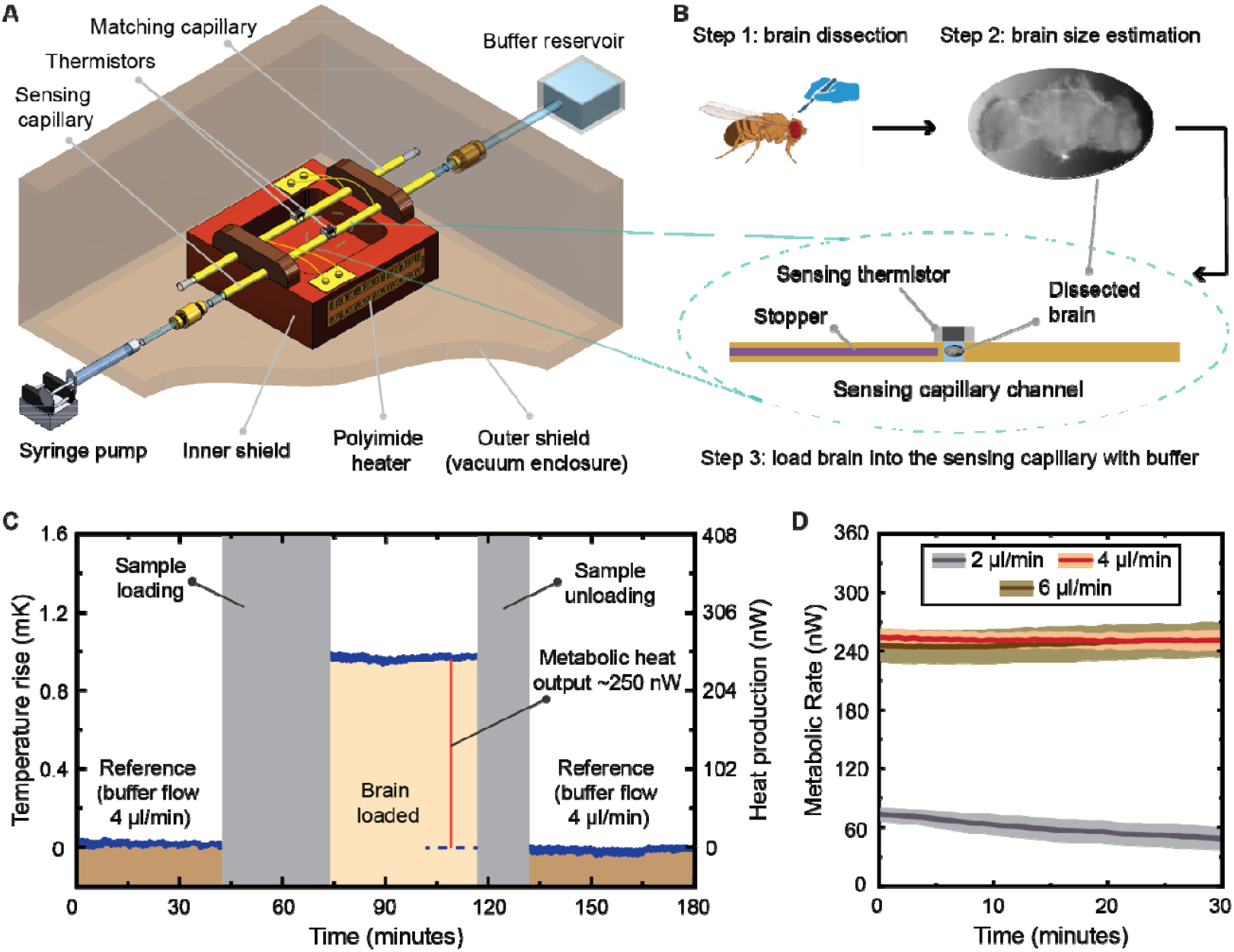
Method for measuring metabolic output in *Drosophila* brains. **(A)** Illustration of the high-resolution biocalorimeter with the thermal shields, the sensing and matching capillary channels, the syringe pump and the buffer reservoir. **(B)** For metabolic output measurements, the brain is dissected from a *Drosophila* fly, imaged for dry mass estimation, and then loaded into the sensing capillary filled with the buffer. The brain is located near the sensing thermistor by the stopper and the measurement begins. **(C)** A trace of the metabolic rate measurement of a brain from a 10-day-old female fruit fly (*y sc v* genotype) with a buffer flow rate of 4 μl/min. The reference temperature is recorded before the measurement. Once the brain is loaded near the thermistor, the metabolic output from the brain raises the temperature in its vicinity, which is recorded by the sensing thermistor. Loading the brain perturbs the system, which takes ∼20 minutes to stabilize. Following stabilization, the measurement is performed for 30 □ 40 minutes, after which the brain is unloaded, and the reference signal is recorded again. The difference in the temperature signal when the brain is in and out of the sensing capillary gives an estimate of the metabolic output of the brain. **(D)** The optimum buffer flow rate is determined to be 4 μl/min for measuring the metabolic activity of *Drosophila* brains. Metabolic output measurements were performed with several brains (10-day-old, female flies, *y sc v* genotype) at buffer flow rates of 2 μl/min, 4 μl/min, and 6 μl/min. The plot shows the mean (solid line) and the standard error of mean (lighter shaded region) of all measured traces over 30 minutes of measurement at these flow rates.

Continuous flow of a physiological buffer solution is necessary for supplying nutrients and oxygen to the explanted brain and maintaining it alive during the calorimetric measurements. We used a buffer made of Schneider’s Drosophila medium supplemented with 1% Antibiotic-Antimycotic solution (Invitrogen), 10% fetal bovine serum (FBS), and 10 μg/ml insulin, which has been previously shown to maintain the health and viability of Drosophila tissues in culture for over 2 days (*31, 32*). The optimum buffer flow rate for the metabolic activity measurements of *Drosophila* brains was found to be 4 μl/min (explained further in the next section). The key calorimetric parameters, i.e. temperature resolution, thermal conductance, and time constant of the sensing capillary were characterized as ∼30 μK (Δ*T*), ∼255 μW/K (*G*_th_) and ∼40 seconds, respectively, with a 4 μl/min buffer flow (fig. S2). The heat output from the brain can be estimated (*33*) from the temperature change near the sensor, given by 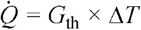. The heat power resolution of the calorimeter was hence determined as ∼7.6 nW with a buffer flow of 4 μl/min.

### Quantification of metabolic activity in *Drosophila* brains

To measure the metabolic heat output from individual fly brains, wild-type flies were synchronized to a 12-hour light/12-hour dark cycle and maintained in an incubator at a constant temperature of 25°C and 50% relative humidity for a minimum of 3 days prior to measurements. Experiments were started two hours after light onset to ensure consistent metabolic activity comparisons across various conditions. To quantify the metabolic heat output of an individual *Drosophila* brain, we dissected a brain using the buffer described earlier, that preserves the health and metabolic activity of dissected tissues for extended periods (*31, 32*). After dissection and before loading it into the calorimeter setup, we optically imaged the brain to trace and measure the largest cross-sectional area of the brain (fig. S6A), which was then raised to the 1.5^th^ power to approximately estimate the brain volume and scaled with a pre-determined volume-to-mass proportionality constant (26.3 μg/mm^3^) to further estimate the dry mass of the brain (see Methods, fig. S6 and fig. S8). The estimated dry mass is used later to determine the normalized metabolic output of the brain from its absolute metabolic output measured via calorimetry.

After imaging, we transferred the brain into the calorimeter’s reservoir and subsequently loaded it into the sensing capillary using the fluid flow generated with a syringe pump (Fig. 1A□B). The brain was held near the sensing thermistor by the stopper and an opposing fluid flow. The buffer flow was then set to the desired continuous flow rate (4 μl/min), after which the calorimeter needed ∼20 minutes for thermal stabilization before beginning the measurement. The measurement was performed for ∼1 hour after loading the brain, and then the brain was unloaded. The temperature changes at the thermistors on the sensing and the matching capillaries were measured using two independent Wheatstone bridge circuits and lock-in amplifiers (SR830, Stanford Research Systems) and the temperature rise caused by the metabolic heat output of the brain was determined as a differential signal between the sensing and the matching capillary sensors to eliminate the common-mode thermal drift. The instrumentation and the measurement scheme are further described in Methods and in fig. S4B. The metabolic heat output of the brain was estimated from: 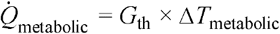.Fig. 1C plots the trace of a complete metabolic rate measurement from the brain of an individual, 10-day-old female fly (*y sc v* genotype), whose metabolic heat output was measured to be ∼250 nW. This trace also shows that the temperature signal remains stable for at least ∼1 hour of continuous measurement with the brain loaded in the calorimeter.

Upon performing measurements with a buffer flow rate of 4 μl/min, the average metabolic output per brain in 10-day-old female flies (*y sc v* genotype) was measured to be ∼256 nW. To investigate the effect of buffer flow rate on the observed metabolic outputs, we performed these measurements at flow rates of 2 μl/min and 6 μl/min. Fig. 1D plots the averaged traces of the metabolic output measured for female brains with 2, 4 and 6 μl/min buffer flow rates. For a flow rate of 2 μl/min, we observed a very low heat output (∼62 nW), that was not stable and gradually kept reducing over time, suggesting that the brains might not be healthy and metabolically active due to an insufficient supply of nutrients and/or oxygen. We observed a stable mean metabolic output of ∼247 nW at a flow rate of 6 μl/min, indistinguishable from the output at 4 μl/min. We also determined the mass-normalized metabolic rate by estimating the dry mass of each brain, following the procedure explained in Methods and in fig. S6, S7. The distribution of the absolute and normalized metabolic rates at the three flow rates (2, 4 and 6 μl/min) are shown in fig. S5A and fig. S5B, respectively. This flow dependency study confirms that a continuous flow of the buffer at 4 μl/min is sufficient to maintain a *Drosophila* brain metabolically active, and all following measurements are performed at the flow rate of 4 μl/min.

### Validation of the metabolic activity measurements

Previous studies using respirometry have demonstrated that cellular metabolism responds dynamically to environmental and pharmacological stimuli, including changes in oxygen availability (*34*), or exposure to mitochondrial inhibitors (*35*). To test whether metabolic heat output of brains is reduced upon exposure to mitochondrial inhibitors, we investigated the effects of disrupting the electron transport chain (ETC) and oxidative phosphorylation. Specifically, we used rotenone, a complex I inhibitor that blocks electron transfer from NADH to ubiquinone, thus reducing ATP production, and antimycin, a complex III inhibitor that prevents electron transfer from ubiquinol to cytochrome c, thereby decreasing ATP synthesis and altering mitochondrial respiration (*36*).

We measured the absolute and normalized metabolic rates of the brains without inhibitor (control), with rotenone (0.5 μM) and with antimycin (0.5 μM and 20 μM). For these experiments, we incubated *Drosophila* brains (from 10-day-old female flies, *y sc v* genotype) in buffer with desired concentrations of rotenone and antimycin for an additional 1 hour at room temperature after dissection and before loading into the calorimeter for measurement (Fig. 2A). For a fair comparison with the experiments performed without the inhibitors, the brains were also incubated for one hour in the control buffer. The metabolic output of brains incubated in the control buffer for 1 hour was identical to that measured in the same buffer without the 1-hour incubation, indicating that a 1-hour incubation in the normal, control buffer does not affect brain metabolism (Fig. 2B). In contrast, exposure to 0.5 μM antimycin resulted in a ∼25% decrease in the metabolic heat output, while 20 μM antimycin led to a ∼80% decrease compared to the control. Similarly, exposure to 0.5 μM rotenone resulted in a ∼60% decrease in the metabolic output. The averaged traces and the distribution of the absolute metabolic rates are plotted in Fig. 2C□D, respectively. Further, we observed that the normalized metabolic rates of brains exposed to antimycin or rotenone were also significantly lower than those of the controls (Fig. 2E). These results showing reduced metabolic heat output following mitochondrial inhibitor treatment are consistent with their known effects on ATP synthesis and demonstrates the sensitivity of our calorimetric method for detecting changes in metabolic activity in individual brains.

**Fig. 2:**
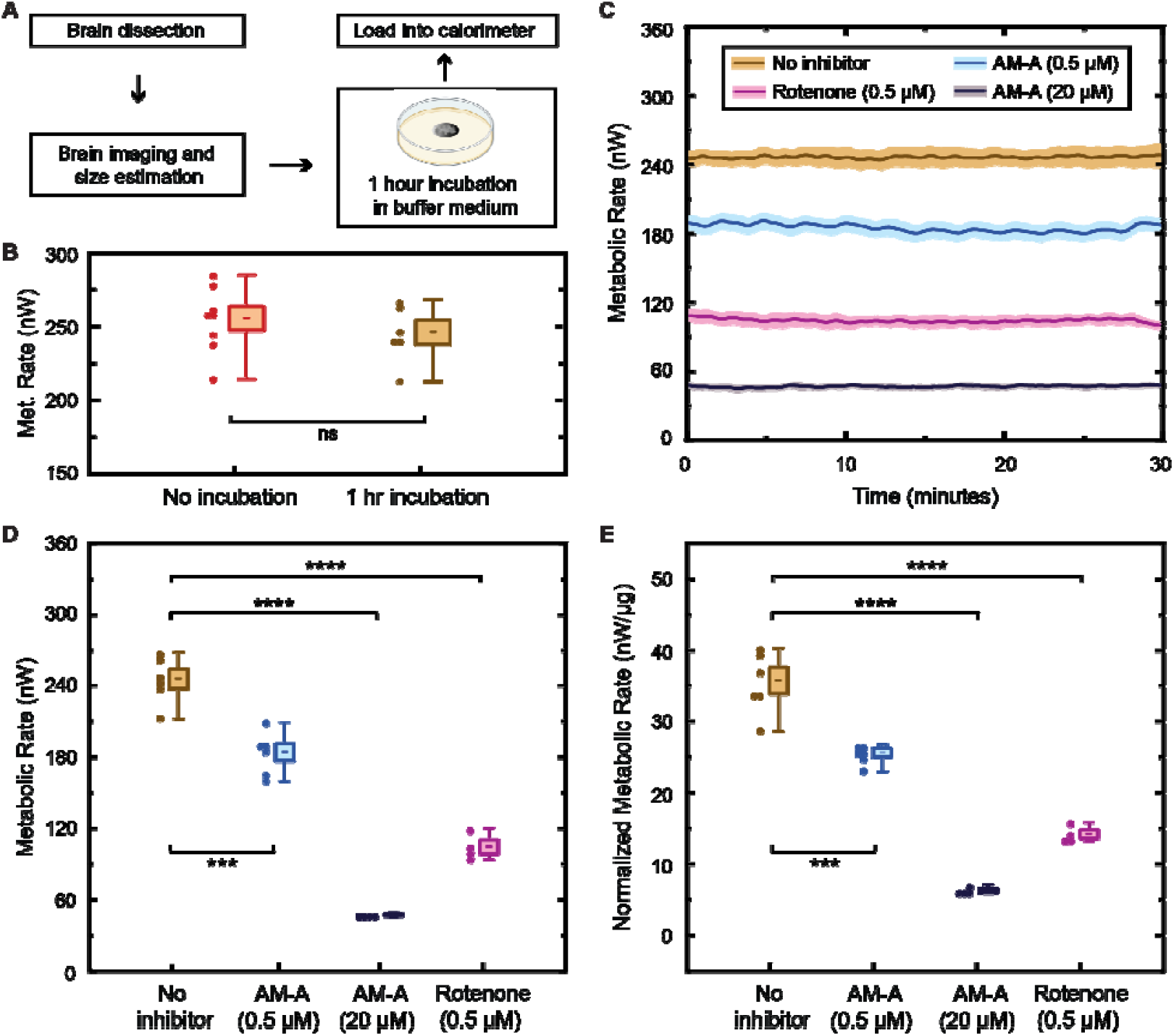
Effect of mitochondrial inhibitors on brain metabolism. **(A)** The method followed for these measurements includes a 1-hour incubation time in the buffer-inhibitor media after imaging and before loading the brains into the calorimeter for measurement. **(B)** Control experiments were performed to show that the additional 1-hour incubation time has no effect on the metabolic rate of brains measured with the control buffer without any inhibitor. **(C)** The average metabolic rate traces of the brains in buffer without any inhibitor, with antimycin-A (AM-A) at 0.5 μM and 20 μM concentrations, and with rotenone at 0.5 μM concentration are plotted, showing the mean (solid line) and the standard error of mean (SEM, lighter shaded region). **(D)** Distribution of the absolute metabolic rates of the brains with the above inhibitors is plotted. A suppression in metabolic rates is observed due to the inhibitors. **(E)** Distribution of the normalized metabolic rates of the brains with the inhibitors is shown, which follows a similar trend. The distribution plots show the individual data points (solid circles), the mean (dash), the SEM (shaded box), and the maximum and minimum values (whiskers). A *p*-value > 0.05 suggests there is no significant (ns) difference between data sets, and □ □ □ □, □ □ □, □ □, and □, indicate *p*□≤ □0.0001, *p*□≤ □0.001, *p* ≤ 0.01, and *p* ≤ 0.05, respectively.

### Quantification of metabolic activity in fly brains across different genders and genotypes

Next, we investigated how brain metabolism in *Drosophila* varies as a function of sex and genotype. Following the workflow outlined in Fig. 1B, we dissected brains, imaged them to estimate size and dry mass, and then loaded them into the calorimeter with the buffer medium to measure the metabolic heat output. Surprisingly, we found that the brains of young (10-day-old) females exhibited significantly higher metabolic heat output than males, in both the *y sc v* and *w*^*1118*^ wild-type genotypes (Figs. 3A–C). To determine whether this difference was due to brain size, we estimated dry mass by imaging each brain, measuring the largest cross-sectional area, and applying this value to our dry mass estimation pipeline (Methods; fig. S7). Our analysis revealed that female and male brains had comparable dry masses (∼6–8 μg; fig. S7). This result was striking because it indicated that the elevated heat output of female brains cannot be explained by differences in brain size. Indeed, even after normalizing metabolic output by dry mass, female brains continued to exhibit significantly higher mass-specific metabolic activity than male brains (Fig. 3D). Taken together, these findings demonstrate a robust sex difference in brain metabolism: young female *Drosophila* brains consistently show higher metabolic activity per unit mass compared to male brains, regardless of genotype. The underlying cellular and molecular mechanisms responsible for this sex-specific metabolic enhancement remain to be elucidated (see Discussion).

**Fig. 3:**
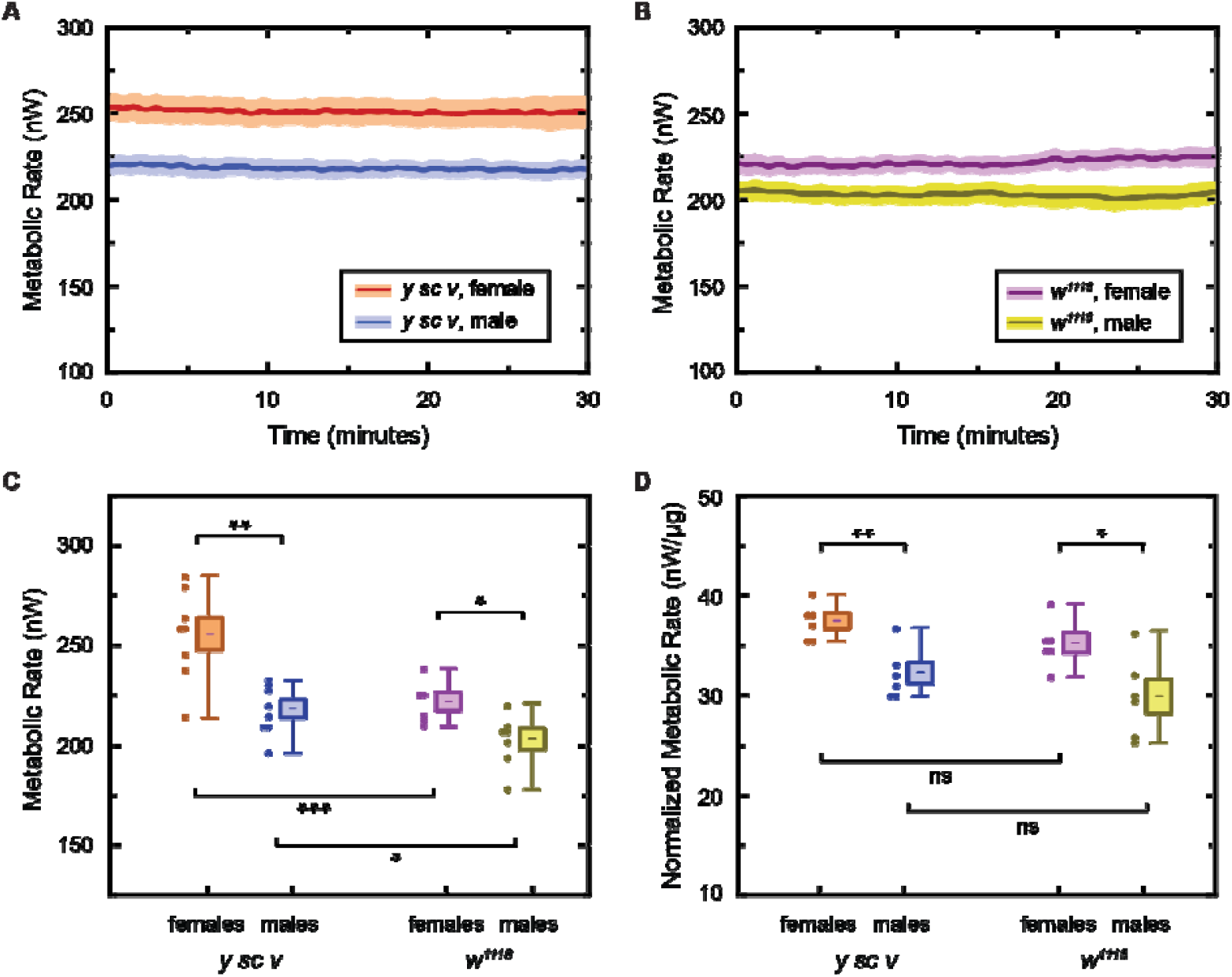
Metabolic activity in fly brains across different genders and genotypes. **(A)** Averaged metabolic output as a function of time is shown for female and male brains of 10-day-old flies from the *y sc v* and **(B)** *w*^*1118*^ genotypes. The mean (solid line) and the standard error of mean (SEM, lighter shaded region) are shown. **(C)** Distribution of absolute metabolic rates of the measured brains is plotted. In both genotypes, females show a higher absolute metabolic rate than males. Both genders of the *y sc v* genotype show a higher absolute metabolic rate compared to the respective genders of the *w*^*1118*^ genotype. **(D)** Distribution of normalized metabolic rates of the measured brains is shown. In both genotypes, females show a higher normalized metabolic rate than males. However, there is no significant difference between the normalized metabolic rates of the same gender across the two genotypes. The distribution plots show each measured data point (solid circles), the mean (dash), the SEM (shaded box), and the maximum and minimum values (whiskers). A *p*-value > 0.05 suggests there is no significant (ns) difference between data sets, and □ □ □ □, □ □ □, □ □, and □, indicate *p*□≤ □0.0001, *p*□≤ □ 0.001, *p* ≤ 0.01, and *p* ≤ 0.05, respectively.

Further, we observed that brains from the *y sc v* genotype exhibited higher metabolic heat output compared to those from the *w*^*1118*^ genotype in both males and females (Fig. 3A-C). Interestingly, *y sc v* brains also tended to have a slightly greater dry mass than *w*^*1118*^ brains, although this difference did not reach statistical significance (fig. S7). This suggests that the elevated heat output in *y sc v* brains may partly reflect their larger overall tissue mass. To account for this, we normalized heat production to brain dry mass. After normalization, metabolic heat output was comparable across genotypes, indicating that the intrinsic metabolic rate per unit mass is not different between *y sc v* and *w*^*1118*^ genotypes (Fig. 3D). Overall, these results provide novel insights into the metabolic dynamics of *Drosophila* brains, highlighting the potential for further studies on metabolic variation due to genetic and physiological factors.

### Quantification of metabolic activity in fly brains across lifespan

Understanding how metabolic activity in specific organs, such as the brain, changes over the lifespan of an organism is crucial due to its potential contribution or correlation to age-related diseases (*5*). To address this, we examined age-related changes in the metabolic activity of *Drosophila* brains in both sexes. We found that female brains exhibited higher metabolic output at a young age, which gradually declined over the lifespan (Fig. 4A, C, E, F). Specifically, metabolic activity decreased substantially between day 10 and day 30, after which values either stabilized or showed a modest increase by day 45. Interestingly, a similar late-life increase in metabolic heat output was observed in females of both genotypes, suggesting that this phenomenon is robust (see Discussion). Additionally, when we quantified normalized metabolic activity, we found it to be significantly higher in younger female flies (day 10) across all genotypes compared to older flies at day 30 or day 45 (Fig. 4G, H), despite comparable dry masses (fig. S9). This indicates a dynamic pattern of metabolic change in female brains over the lifespan.

**Fig. 4:**
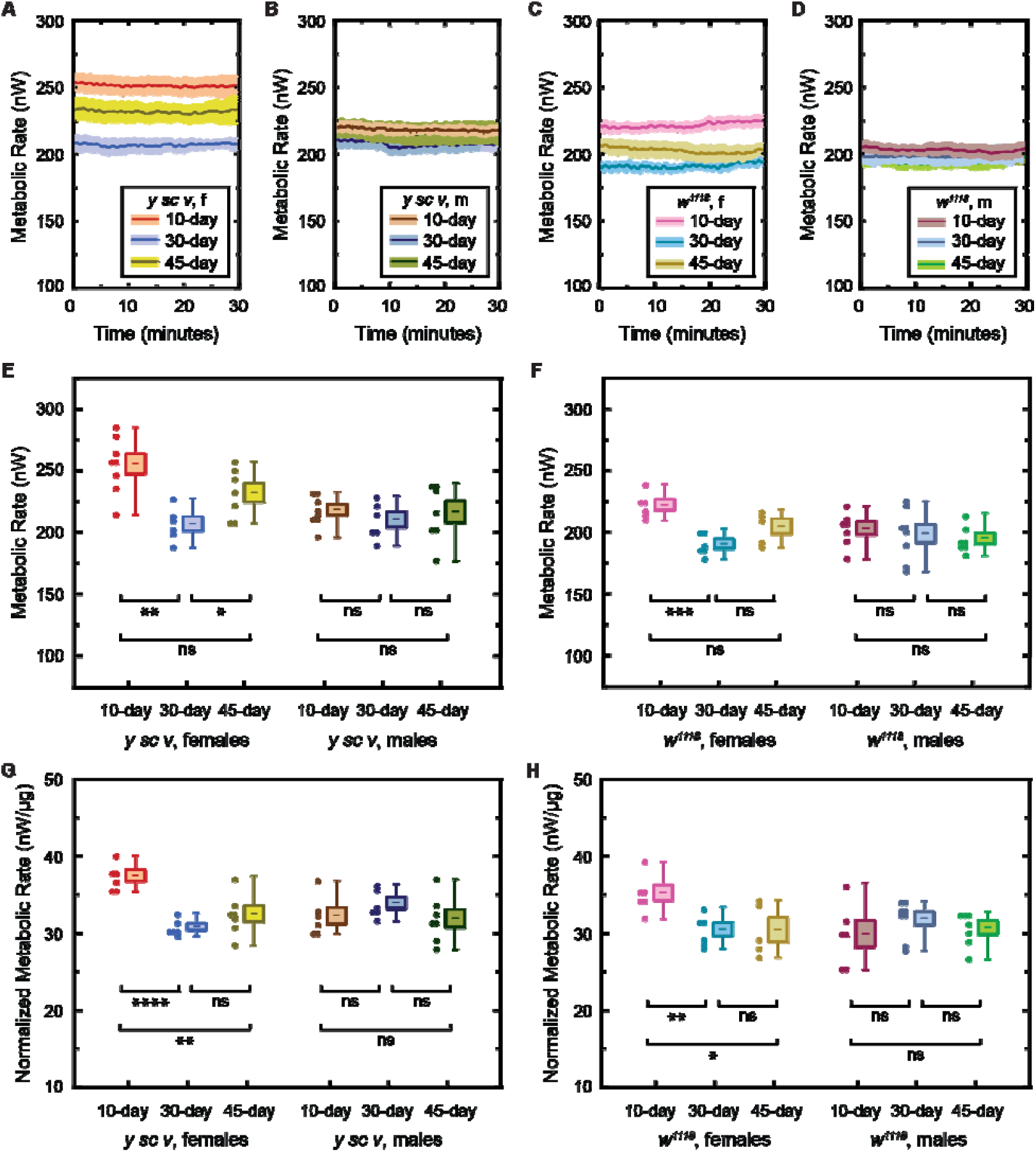
Metabolic activity in fly brains across lifespan. **(A)** Averaged absolute metabolic output as a function of time is shown for brains of 10-, 30- and 45-day-old female flies of the *y sc v* genotype (*y sc v*, f), **(B)** male flies of the *y sc v* genotype (*y sc v*, m), **(C)** female flies of the *w*^*1118*^ genotype (*w*^*1118*^, f) and **(D)** male flies of the *w*^*1118*^ genotype (*w*^*1118*^, m). The mean (solid line) and the standard error of mean (SEM, lighter shaded region) are shown. **(E)** Distribution of absolute metabolic rates of the measured brains from female and male flies of the *y sc v* and **(F)** *w*^*1118*^ genotypes. The absolute metabolic rates of the female brains sharply reduce from ages day 10 to day 30, and then slightly increase or remain constant from day 30 to day 45. In contrast, the absolute metabolic rates of the male brains remain constant throughout their lifespan, in both the genotypes. **(G)** Distribution of normalized metabolic rates of the measured brains from female and male flies of the *y sc v* and **(H)** *w*^*1118*^ genotypes. In both genotypes, the normalized metabolic rates of the female brains are higher at day 10 compared to day 30 or day 45, while the normalized metabolism in males remained constant throughout their lifespan. The distribution plots show each measured data point (solid circles), the mean (dash), the SEM (shaded box), and the maximum and minimum values (whiskers). A *p*-value > 0.05 suggests there is no significant (ns) difference between data sets, and □ □ □ □, □ □ □, □ □, and □, indicate *p*□≤ □0.0001, *p*□≤ □0.001, *p* ≤ 0.01, and *p* ≤ 0.05, respectively.

By contrast, males showed relatively stable absolute metabolic rates (Fig. 4B, D, E, F) as well as normalized metabolic rates (Fig. 4G, H) from day 10 to day 45 in both *y sc v* and *w*^*1118*^ genotypes. Together, these findings provide the first quantitative description of brain metabolic aging in *Drosophila*, revealing distinct sex-specific trajectories: females exhibit an early-life peak followed by decline, whereas males maintain a more constant metabolic profile.

### Quantification of metabolic activity in mitochondrial mutant fly brains

We next investigated how mutations in mitochondrial quality control genes such as *Parkin*, which are linked to Parkinson’s disease, affect the metabolic activity in brains. *Parkin*, encoded by the *PARK2* gene, is vital for essential for maintaining mitochondrial quality control by functioning as an E3 ubiquitin ligase that tags damaged mitochondria for degradation via the mitophagy pathway (*37, 38*). Disruption of this pathway due to *Parkin* mutations leads to the accumulation of dysfunctional mitochondria, impaired energy production, increased oxidative stress, and ultimately neuronal dysfunction. Consistent with this, *Parkin* null *Drosophila* exhibit Parkinsonian-like phenotypes, including reduced lifespan, motor deficits, sterility, mitochondrial abnormalities, and dopaminergic neurodegeneration (*39*).

In our study, we assessed the consequences of Parkin-mediated mitochondrial dysfunction on brain metabolism using *Drosophila* carrying heterozygous (*park*^*1*^*/+*) or homozygous (*park*^*1*^) mutations. Brains from heterozygous mutants exhibited a mean metabolic output of ∼246 nW, whereas brains from homozygous mutants showed a reduced output of ∼211 nW, corresponding to an ∼15% decline (Fig. 5). Notably, this reduction remained evident after normalization to brain dry mass, indicating that the observed effect is not attributable to differences in brain size but rather reflects a decrease in metabolic activity. These findings demonstrate that loss of Parkin function leads to a significant reduction in brain metabolic activity, consistent with impaired mitochondrial performance. The data provide direct functional evidence that Parkin-mediated mitochondrial quality control is required to sustain normal metabolic output in the brain, and disruption of this pathway results in energy deficits that may contribute to neurodegenerative phenotypes.

**Fig. 5:**
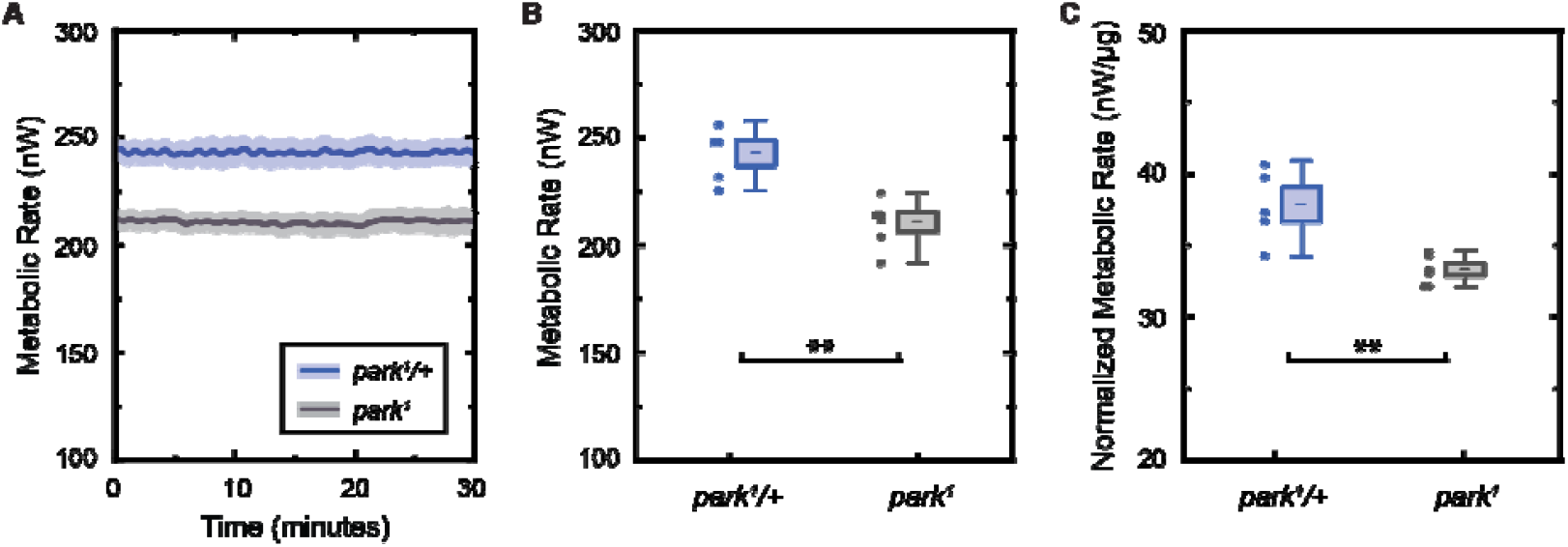
Metabolic activity in brains of mitochondrial mutant flies. **(A)** Averaged absolute metabolic output as a function of time is plotted for heterozygous (*park*^*1*^*/+*) and homozygous (*park*^*1*^) mutant fly brains. The mean (solid line) and the standard error of mean (SEM, lighter shaded region) are shown. **(B)** Distribution of the absolute metabolic rates is plotted, showing a ∼15% suppression in the metabolism of *park*^*1*^ mutant brains compared to control brains. **(C)** Distribution of the normalized metabolic rates also shows a similar reduction in metabolism of the homozygous mutant brains. The distribution plots show each measured data point (solid circles), the mean (dash), the SEM (shaded box), and the maximum and minimum values (whiskers). A *p*-value > 0.05 suggests there is no significant (ns) difference between data sets, and □ □ □ □, □ □ □, □ □, and □, indicate *p*□≤ □0.0001, *p*□≤ □0.001, *p* ≤ 0.01, and *p* ≤ 0.05, respectively.

### Quantification of metabolic activity in other *Drosophila* tissues

Finally, we performed measurements quantifying metabolic activity in other *Drosophila* tissues like ovaries and testes, demonstrating the general capabilities of the developed calorimetric method. In these studies, we followed similar dissection, loading, and measurement protocols as described in Fig. 1. One representative measurement trace each for an individual ovary and an individual testis are shown in fig. S9A and fig. S9B, respectively. Measurements performed on 6 ovaries and 6 testes revealed an average metabolic heat output of ∼132 nW per ovary and ∼83 nW per testis, in 10-day-old flies of the *y sc v* genotype (Fig. 6A□B).

**Fig. 6:**
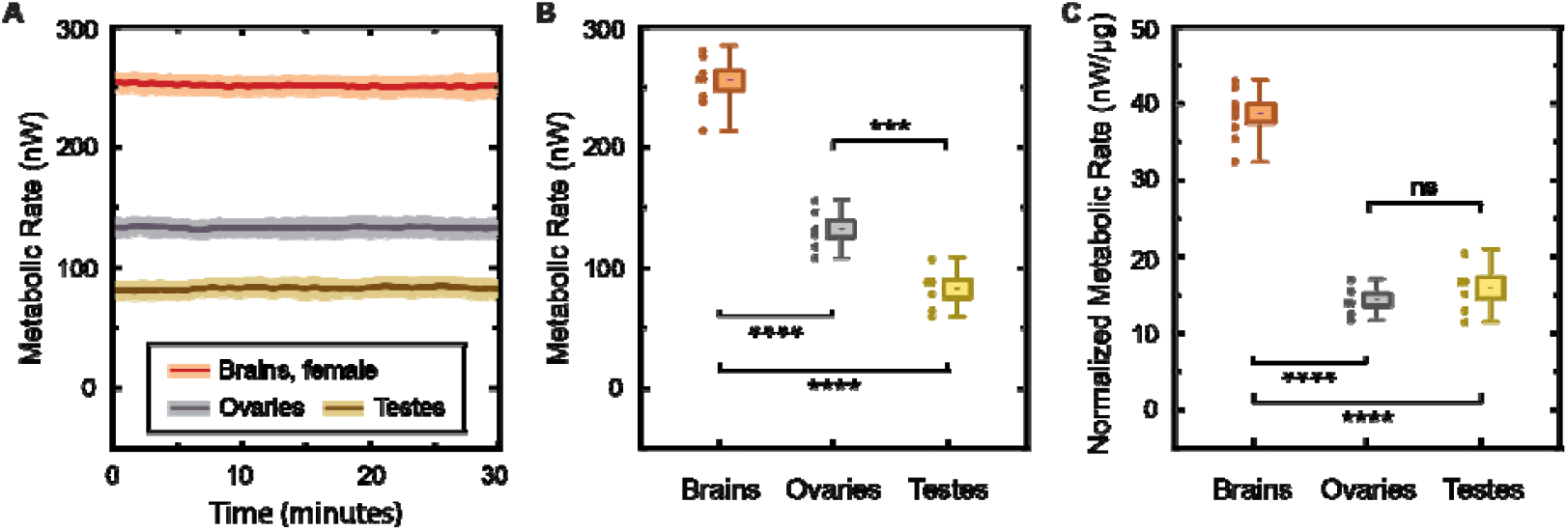
Metabolic activity of other *Drosophila* tissues. **(A)** Averaged absolute metabolic output as a function of time for female brains, ovaries, and testes of 10-day-old flies (*y sc v* genotype). The mean (solid line) and the standard error of mean (SEM, lighter shaded region) are shown. **(B)** Distribution of the absolute metabolic rates of the measured tissues. The mean metabolic output of the female brains, ovaries, and testes were ∼256 nW, ∼132 nW, and ∼83 nW, respectively. **(C)** Distribution of normalized metabolic rates of the brains, ovaries and testes. The mean of the normalized metabolic outputs of the female brains, ovaries, and testes were ∼38.7 nW/μg, ∼14.4 nW/μg, and 15.9 nW/μg, respectively. The normalized metabolic outputs of the ovaries and the testes are similar (∼15 nW/μg), while that of the brains is ∼2.5-fold higher. The distribution plots show each measured data point (solid circles), the mean (dash), the SEM (shaded box), and the maximum and minimum values (whiskers). A *p*-value > 0.05 suggests there is no significant (ns) difference between data sets, and □ □ □ □, □ □ □, □ □, and □, indicate *p*□≤ □0.0001, *p*□≤ □0.001, *p* ≤ 0.01, and *p* ≤ 0.05, respectively.

To compare normalized metabolic rates across tissues, we measured dry mass with an ultra-microbalance (see Methods and fig. S8A). The mean dry masses of female brains, ovaries, and testes were found to be ∼6.6 μg, ∼9.2 μg, and ∼5.2 μg, respectively (fig. S9B). Thus, we estimated the normalized metabolic outputs to be ∼38.7 nW/μg for female brains, ∼14.4 nW/μg for ovaries, and ∼15.9 nW/μg for testes, in 10-day-old flies of the *y sc v* genotype (Fig. 6C). Our observations indicate that the brain’s normalized metabolic activity is significantly (∼2.5 fold) higher than that of ovaries and testes, demonstrating that the brain is one of the most energy-consuming organs. Our demonstrations underscore the versatility and efficacy of this calorimetric method in providing detailed metabolic insights across different tissues in addition to brains, establishing it as a powerful tool for metabolic studies in a range of small biological tissues.

## DISCUSSION

We present a novel calorimetric tool that enables, for the first time, direct, real-time measurements of metabolic output from small neural tissues, such as live explanted *Drosophila melanogaster* brains. This advance, in contrast to past work on probing cellular temperatures (*40*) or metabolism of individual model organisms (*24, 25*), allows for the first time a direct approach to monitoring the metabolism of live tissues of model organisms. The developed biocalorimeter can detect power output changes as small as ∼7.6 nW with a response time of ∼40 seconds, while preserving viability of brains or other tissues (ovaries and testes) via continuous buffer perfusion (∼4 μl/min), enabling stable metabolic measurements on individual explanted brains for over 1 hour. Our experiments directly illustrate the calorimeter’s sensitivity in detecting metabolic shifts. Exposure to mitochondrial inhibitors (0.5 μM antimycin: ∼25% reduction; 20 μM antimycin: ∼80% reduction; 0.5 μM rotenone: ∼60% reduction) reduced brain metabolic activity, consistent with their known disruption of ATP synthesis via the electron transport chain. These results demonstrate the ability of our tool to quantify dynamic metabolic responses to pharmacological and environmental stimuli in individual, intact brains.

Our quantification of metabolic activity across genders and genotypes uncovered novel insights. Female *Drosophila* brains exhibited significantly higher metabolic rates than male brains for both *y sc v* and *w*^*1118*^ genotypes even when normalized for dry mass. This disparity in metabolic output cannot be solely attributed to the cell numbers between genders, as recent studies have not shown significant differences in the cell numbers between male and female *Drosophila* brains (*41*). Potential factors that could contribute to higher metabolic rates in female brains include increased neuronal activity, hormonal influences on metabolism, or genetic variations affecting metabolic pathways.

Age-related analyses revealed distinct metabolic trajectories: female brains showed a decline in metabolic rate from day 10 to day 30, which stabilized or slightly increased by day 45, while male brains remained stable throughout day 10 to day 45. The basis of this late-life metabolic rise remains unclear, but several possibilities exist. It may reflect compensatory upregulation of energy metabolism in aging neurons or glia, an increased contribution of non-neuronal cell types, or a stress-induced elevation in mitochondrial activity. Alternatively, it could arise from age-related changes in tissue composition or increased cell turnover. Regardless of the underlying cause, these results highlight that brain metabolism does not decline linearly with age but instead exhibits distinct phases, with an early-life decrease followed by stabilization or partial recovery in later life. Additionally, th ese gender-specific patterns, previously unreported, highlight the capability to track metabolic aging in brains, providing insights into the metabolic basis of age-related diseases.

In *Drosophila* models of Parkinson’s disease (*39*), homozygous mutants (*park*^*1*^) exhibited a ∼15% reduction in brain metabolic output (∼211 nW) compared to heterozygous mutants (*park*^*1*^*/+*, ∼246 nW), even after dry mass normalization. These findings demonstrate that mitochondrial dysfunction caused by Parkin deficiency leads to lower metabolic outputs. These studies, for the first time, show direct quantitative tracking of neurodegenerative disease-relevant metabolic alterations in brains.

Finally, we demonstrated the capability to quantify metabolism in other tissues of *Drosophila* beyond brains, such as reproductive organs like ovaries and testes. We reported metabolic outputs of ∼132 nW and ∼83 nW in ovaries and testes of 10-day-old *Drosophila*, compared to the ∼256 nW output in brains of female 10-day-old Drosophila. Notably, the brain’s normalized metabolic rate (∼38.7 nW/μg) was significantly higher compared to ovaries (∼14.4 nW/μg) and testes (∼15.9 nW/μg), demonstrating that the brain is a highly energy-intensive organ.

In conclusion, the presented calorimetric approach represents a transformative tool for probing bioenergetics, enabling precise, real-time metabolic measurements in individual, intact brains of small model organisms like *Drosophila*, providing new insights into brain energetics, aging, and disease. Using this platform, we discovered novel sex-, age-, genotype-, and disease-dependent patterns in brain metabolism. By bridging the gap in quantitative bioenergetics, our studies open new avenues for understanding cellular energy dynamics in health and disease. Further, this method can be extended beyond *Drosophila* brains to organoids and drug-screening platforms, paving the way for widespread application in small-tissue and precision medicine research.

## MATERIALS AND METHODS

### Calorimeter design and fabrication

Two cylindrical capillary tubes (0.75 mm and 1 mm inner and outer diameters, respectively) made of Teflon PFA are prepared as the sensing and the matching capillary sensors (Fig. 1A and fig. S1A). The process for the fabrication and assembly of the sensing capillary is shown by Steps 1-7 in fig. S1B. The PFA capillary is cleaned with acetone and iso-propyl alcohol to prepare for metal deposition (Step 1). A 5/100 nm-thin titanium/gold film is deposited using e-beam evaporation throughout the cylindrical surface of the capillary except at a small window at the center (Step 2). This window electrically isolates the metal on the two sides of the capillary and also allows for optical access. A miniature thermistor (Ametherm, NTC, SM06103395), used as the thermometric sensor for metabolic measurements, is mounted at the center along the length of the capillary, soldered to the gold layers on either side such that the gold layers extend the electrical connections from the two terminals of the thermistor to the external electrical instrumentation (Step 3).

The outer shield (OS, 12×12×8 cm^3^) is a vacuum enclosure while the inner shield (IS, 7×7×3.5 cm^3^) is U-shaped and is enclosed on 5 sides but open on one side, and both shields are made of copper. The sensing capillary with the mounted thermistor is assembled onto the IS, such that ∼20 mm length of the capillary is suspended on the open side of the IS (Step 4). A small copper lid (part of the IS), with a cylindrical groove (1 mm diameter) lined with a thin, soft graphite sheet, encloses the capillary from the top side and is mechanically screwed down to the rest of the IS. The graphite fills the air pockets between the capillary and the groove of the copper lid, and ensures that the capillary is in excellent thermal contact with the IS. The IS is housed inside the OS (Step 5), and the sensing capillary extends out of the OS through hermetically sealed PEEK feedthroughs and connectors (Idex MicroTight series). A multimode optical fiber (Thor Labs, UM22-300, 370 μm outer diameter) is inserted into the capillary from one side and extends till the thermistor on the capillary (Step 6). It acts as a stopper to localize biological samples for metabolic measurements and also acts as a light source for optical imaging of the samples. One end of the capillary (same end through which the stopper is inserted) is connected to a syringe pump, while the other end of the capillary sinks in a reservoir, which together completes a microfluidic system. A physiological buffer is then filled into the capillary for biocalorimetry measurements (Step 7).

The primary difference between the sensing and matching capillaries is that the latter does not extend out of the OS, does not have a stopper, and is not filled with buffer. The process for the fabrication and assembly of the matching capillary follows Steps 1-4 shown in fig. S1B, i.e. it is identical to the process for the sensing capillary till after its assembly with the IS.

### Temperature stabilization

A nested two-shield system is designed to isolate the sensing and matching capillaries from ambient temperature fluctuations. OS and IS are made of highly thermally conductive copper, have large masses to dampen high-frequency temperature oscillations, and are isolated from each other using weak thermal links. The IS is mounted on the OS using four borosilicate spheres (fig. S1A) sitting between conical grooves machined on the IS and the OS, connecting the two shields conductively at approximate line contacts. Vacuum conditions < 10 μTorr are maintained inside the OS, which significantly attenuate the convective heat transfer (*42*). Radiative heat transfer between the shields is reduced by using reflective aluminum sheets (emissivity < 0.07(*28*)) on the shield surfaces. Finally, the OS is externally enclosed with reflective foam insulation to prevent air currents from perturbing the system.

High-precision temperature sensing and active PID-based temperature control are employed to eliminate low-frequency (long-term) temperature drifts in the shields. The instrumentation developed for the temperature feedback control is shown in fig. S4A. AC-driven Wheatstone bridge circuits are used for resistance thermometry(*43*), with precision thermistors for temperature sensing (US Sensor Corp., USP 12838, 10 kΩ nominal resistance at 25°C, –4.4%/K *TCR* (temperature coefficient of resistance)) integrated into the walls of the OS and IS. A temperature change in the shield leads to a change in the resistance of the thermistor, which is detected using a sinusoidal sensing current in the Wheatstone bridge circuit. The output is amplified using an instrumentation amplifier (AD 524), measured as *V*_sens,shield_ (Eq. 1) and provided as an input to the PID control algorithm, which is implemented through LABVIEW. The PID control algorithm, following the general PID control techniques given in Ref. (*44*), compares *V*_sens,shield_ to a pre-determined voltage setpoint *V*_set_ and outputs a feedback voltage *V*_PID_ in the range of –10 V to 10 V. *V*_PID_ is provided as an input to a current-source circuit that uses an NPN transistor (ST Microelectronics 2ST31A) to adjust the heating current supplied to the polyimide heaters (Omega KH series) mounted on the surface of the shields, thus controlling the temperature of the shields around a desired setpoint with high precision. Using the above method, we achieved a temperature stability within ±1 μK in the IS over 10 hours, measured at a 1 mHz bandwidth (∼100 dB attenuation in temperature fluctuations from the ambient to the IS). Our past work(*30*) shows these results and describes the stabilization method and instrumentation in greater detail.

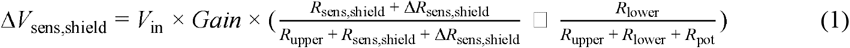

### Thermometry to quantify metabolic heat output

The biological sample is located near the thermistor mounted on the sensing capillary. The measured metabolic heat output is given by 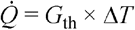, where Δ*T* is the temperature change sensed by the thermistor and *G*_th_ is the thermal conductance of the capillary to the IS. The ability to measure small metabolic heat outputs relies on a high thermometric resolution and a small *G*_th_. To achieve high thermometric resolution, we implement resistance thermometry(*43*) using a high *TCR* miniature thermistor, precise AC-driven Wheatstone bridge circuitry, and common-mode cancellation of thermal drift. The sensing capillary with the integrated thermistor connected to the electrical instrumentation is illustrated in fig. S4B. The sensing thermistor (Ametherm, NTC, SM06103395), connected to the lower right leg of the Wheatstone bridge, has a high *TCR* (−3.95%/K) and a nominal resistance of 10 kΩ (±1%) at 25°C (*R*_sens cap_). A fixed 10 kΩ resistor (*R*_lower_) is connected to the lower left leg of the bridge. The upper right leg consists of a 40 kΩ fixed resistor (*R*_upper_)while the upper left leg consists of a 40 kΩ fixed resistor (*R*_upper_) along with a potentiometer (*R*_pot_, 0 □ 500 Ω) for fine balancing of the bridge. All fixed resistors in the bridge are precision, ultra-low *TCR* resistors (Vishay Z201 Series Z-Foil, ±0.2 ppm/K), and the potentiometer is from Vishay Spectrol 534 series with a 20 ppm/K *TCR*. The temperature change due to the metabolic output from a sample causes a change in the DC resistance of the thermistor, which is detected using an AC sensing voltage (*V*_AC_).

Using a modulated sensing current helps attenuate the voltage noise introduced in the sensor circuits through 1/*f* noise sources and other DC offsets. Toward this, the bridge is excited using a 1 V peak-to-peak sinusoidal voltage (*V*_AC_) at a frequency in the range of 20 □ 40 Hz (*f*_AC_) using an Agilent 33120A waveform generator. The change in voltage across the thermistor on the sensing capillary is output from the Wheatstone bridge (at *f*_AC_) and amplified by an AD 524 instrumentation amplifier. This voltage, i.e. Δ*V*_sens cap_, given by Eq. 2, is measured in a ∼16 mHz bandwidth using a Lock-in mechanism (SRS 830) and recorded using a NI PCI-6014 DAQ card. Δ*V*_sens cap_ is converted to the corresponding resistance change (Δ*R*_sens cap_) and temperature change (Δ*T*_sens cap_) using Eqns. 3-4.

A matching capillary, identical to the sensing capillary but without a biological sample, is designed to implement a common-mode cancellation scheme to eliminate long-term temperature drifts. The temperature change in the matching capillary is measured following similar circuitry and instrumentation as for the sensing capillary, described above. Temperature change in the matching capillary is primarily caused by ambient temperature fluctuations. The temperature change caused due to the metabolism of the biological sample is thus estimated by subtracting the temperature change in the sensing capillary (Δ*T*_sens cap_) and the temperature change in the matching capillary (Δ*T*_mat cap_). The measured temperature change of the biological sample is converted to the metabolic heat output 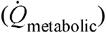 using the pre-calibrated *G*_th_ (Eq. 5).

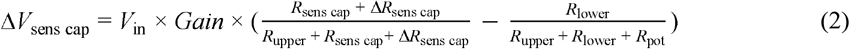

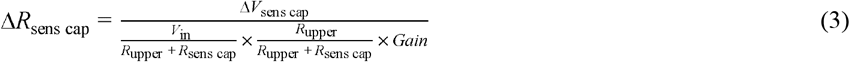

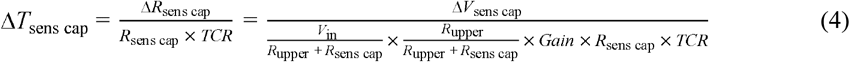

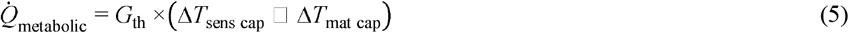

### Thermal conductance, heat power resolution, and time constant

Characterizing the thermal conductance of the suspended calorimeter requires applying a DC voltage offset (*V*_DC, off_) in addition to the AC sensing voltage (*V*_AC_). *V*_DC,off_ causes a known power dissipation on the thermistor due to Joule heating, leading to a constant temperature rise that is measured using *V*_AC_ and the measurement scheme described in the previous section. The slope of the power dissipated to the temperature rise, corresponds to the thermal conductance (*G*_th_). The *G*_th_ of the sensing capillary calorimeter is measured (shown in fig. S2A) in the presence of a 4 μl/min buffer flow through the capillary. In this measurement, DC power was dissipated in the calorimeter from 0 □ 160 nW in steps of 20 nW, and the corresponding temperature rise was measured from ∼0 □ 630 μK, resulting in a *G*_th_ of ∼255 μW/K. As shown in fig. S3A, we also measured the *G*_th_ with buffer flow rates of 0 μl/min, 2 μl/min, and 6 μl/min, and found them to be ∼210 μW/K, 228 μW/K, and 290 μW/K, respectively.

The thermal time constant of the calorimeter was characterized as ∼40 seconds with a 4 μl/min buffer flow, measured at a ∼16 mHz bandwidth (fig. S2B), the same bandwidth as used for the metabolic heat output measurements. To measure this, power was dissipated from 0 □ 200 nW in steps of 100 nW. The time taken by the calorimeter to reach ∼63.2% of its final temperature change due to the step power input, is the thermal time constant of the calorimeter.

To characterize the heat power resolution, the DC power dissipated was applied from 1.5-18 nW in small steps of 1.5 □ 3 nW. The inset in fig. S2A plots the measured temperature rise at these dissipated powers, showing the temperature measurement noise floor to be ∼30 μK corresponding to a power dissipation of ∼7.6 nW. Hence, the temperature resolution and the heat power resolution of this calorimeter are ∼30 μK and ∼7.6 nW, respectively with a buffer flow rate of 4 μl/min. To validate this further, we applied DC power dissipation in steps of 7.6 nW and could clearly resolve the resulting steps of ∼30 μK in the temperature rise (fig. S3B). Further, we also measured the temperature resolution and the heat power resolution when there is no buffer flow and measured them to be ∼9.5 μK and ∼2 nW, respectively (fig. S3B). While the *Drosophila* brains and tissues studied here need a buffer flow of 4 μl/min for which the heat power resolution is ∼7.6 nW, this calorimeter can offer a higher sensitivity: up to ∼2 nW with no buffer flow. This outlines the ultimate heat sensing capabilities of the developed calorimeter, which can be exploited when employing it for studying metabolic activity in biological samples requiring lower buffer flow rates during measurements. All above measurements are performed at a ∼16 mHz bandwidth.

### Fly husbandry

Adult male and female *Drosophila melanogaster* were entrained under standard 12-hour/12-hour light/dark (LD) cycles at 25°C and 50% relative humidity for a minimum of 3 days prior to dissection. The experiments were conducted at the same time every day to control for the time-of-day variations. The flies used in the study, *y sc v, w*^*1118*^, and *park*^*1*^ were obtained from the Bloomington Drosophila Stock Center.

### Dissection and culture of *Drosophila* brain explants and other tissues

Flies were anesthetized on ice for 5□10 seconds before dissection. Dissections were performed in a sterile petri dish containing an ice-cold modified culture medium (buffer) adapted from the work by Ayaz et al. (*31*), consisting of Schneider’s Drosophila Medium (Invitrogen, Carlsbad, CA) supplemented with 1% Antibiotic-Antimycotic solution (Invitrogen), 10% fetal bovine serum (FBS), and 10 μg/ml insulin. Prior studies have confirmed that these brain cultures maintain identifiable morphological features and neuronal functionality for up to 6 days (*32*). Whole brains, including intact optic lobes, were carefully dissected, with cuticles and trachea removed. Testes and ovaries were similarly dissected in the same ice-cold culture medium, ensuring removal of extraneous tissues as needed. Dissected brains and tissues were rinsed three times in the medium before loading into the calorimeter, which was filled with the same buffer medium at room temperature for the calorimetric measurements. For specific experiments involving mitochondrial inhibition in brains, the following pharmacological agents were added to the culture medium: rotenone (0.5 μM), and antimycin A (0.5 μM and 20 μM).

### Estimation of brain dry mass by imaging

For comparing the mass-specific metabolic outputs of the brains, it was challenging and unfeasible to directly measure the dry mass of each individual brain whose metabolic activity was quantified. Hence, we estimated the mass of the brains by imaging the brains and measuring their largest area of cross-section. The traced area of cross-section of a brain is shown in fig. S6A. The dry mass of a brain is directly proportional to its volume, which can be approximated as the 1.5^th^ power of the cross-sectional area, assuming an isometric scaling. To estimate the proportionality constant relating brain volume to the dry mass, we simultaneously measured the dry mass and the cross-sectional area for 6 female brains of 10-day-old Drosophila flies (*y sc v* genotype). The proportionality constant was estimated to be ∼26.3 μg/mm^3^ by plotting the dry masses against the 1.5^th^ power of the cross-sectional area measured for these 6 brains (shown in fig. S6B).

### Dry mass measurements using ultra-microbalance

A Mettler Toledo XP2U ultra-microbalance (resolution: 0.2 μg) was used for measuring the dry mass of brains, ovaries, and testes (the followed procedure is shown in fig. S8A). Weighing boats (∼1 mm × 1 mm in size) were prepared out of aluminum foil and their masses were measured with the microbalance. The average mass of an empty boat was ∼100 μg. For dry mass measurement, a tissue was first dissected in the buffer medium. Next, it was quickly rinsed (for 2□3 seconds) in de-ionized (DI) water to get rid of the buffer salts and then transferred to a boat. The boat with the tissue was transferred to a temperature-controlled chamber (TestEquity Model TEC1) and heated at 50°C for 5 hours to dehydrate the tissue. Finally, the dry mass of the tissue in the boat was measured using the microbalance and the pre-calibrated mass of the boat was subtracted to obtain the tissue dry mass. The measured dry masses of several female brains, ovaries, and testes from 10-day-old *Drosophila* flies (*y sc v* genotype) are plotted in fig. S8B.

### Data analysis

Basal metabolic rate was calculated for an individual fly brain or tissue by averaging its heat output over ∼30 minutes of stable measurement, as shown in the individual measurement traces in Fig. 1C and fig. S9. Statistical analysis to compare the effects of inhibitors, gender, genotype, age, and disease on metabolic rates, was performed using the two-tailed *t* test in OriginPro 2022 and computing the *p*-values. In the data shown, *p*□≤ □0.0001, *p*□≤ □0.001, *p* ≤ 0.01, and *p* ≤ 0.05 are indicated by □ □ □ □, □ □ □, □ □, and □, respectively. A *p*-value > 0.05 suggests there is no significant (ns) difference between data sets.

## Acknowledgements

We thank the members of the Reddy, Meyhofer, and Yadlapalli laboratories for helpful discussions. We thank the Bloomington Drosophila Stock Center (P40 OD018537) for providing fly strains.

## Funding

This work was supported by National Institutes of Health NIGMS R35GM133737 grant to SY, the Alfred P. Sloan fellowship, the McKnight scholar grant, and the Chan Zuckerberg collaborative pairs grant to SY.

## Author contributions

PR, EM, and SY conceived the work. K P and RM developed the instrument. KP performed all calorimetric experiments. RM assisted with preliminary calorimetric experiments. KP and AB performed all dry mass measurements. QC and SY performed fly husbandry and all dissections. The manuscript was written by KP, PR, EM, and SY with comments and input from all authors.

## Competing interests

The authors declare that they have no competing interests.

## Data and materials availability

All data needed to evaluate the conclusions are present in the paper and/or the Supplementary Materials.

## Notes

### Competing Interest Statement

The authors have declared no competing interest.

### Summary of Updates

Quantitative insights into brain metabolism are essential for advancing our understanding of energy dynamics in the brain. However, current approaches for tracking brain metabolism, metabolic profiling and respirometry, provide only static snapshots of metabolite levels or lack the required resolution. Here, we develop a novel nanowatt-resolution biocalorimeter capable of real-time continuous measurements of heat output to quantitatively measure the metabolism of individual live Drosophila melanogaster brains and investigate how sex, genotype, age, and disease affect brain metabolism. We show for the first time that female brains, across multiple wild-type genotypes, exhibit a significantly higher metabolic rate (~10%) than male brains at a young age (<10 days old) and follow distinct metabolic trajectories across the lifespan. We also find that parkin mutants, a genetic model for Parkinsons disease, exhibit a ~15% reduction in brain metabolic output relative to controls, revealing that defective mitophagy due to parkin deficiency affects brain metabolism. Furthermore, we measure the metabolic rate of reproductive tissues of Drosophila, highlighting the broad applicability of our biocalorimeter. Together, these advances open new avenues for investigating how tissue-specific metabolism is impacted by aging, neurodegeneration, and disease states.

