## Supplementary material for "Direct quantification of the metabolic heat output of individual *Drosophila* brains": supp figures

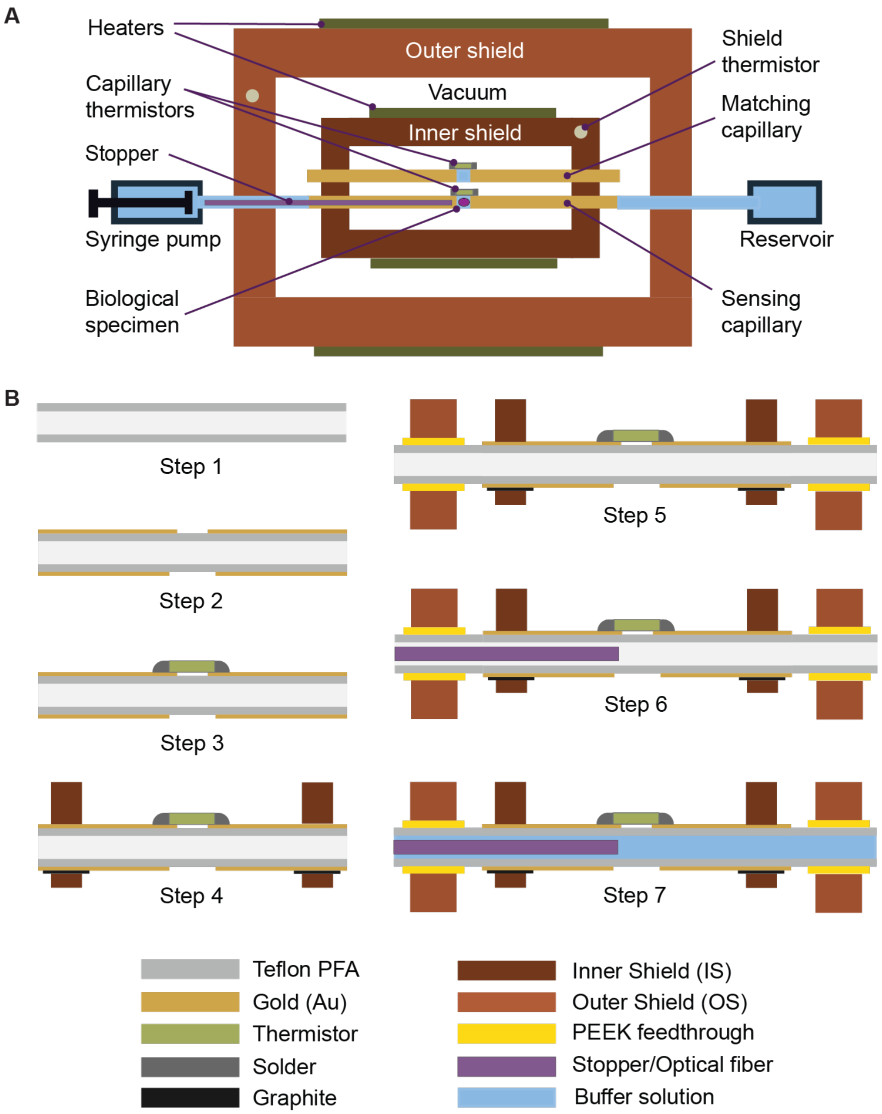


**Fig. S1: Design and fabrication of the calorimeter. (A)** Schematic top view of the developed biocalorimeter**. (B)** Process flow for the fabrication and assembly of the biocalorimeter. Step 1: A Teflon PFA capillary tube is cleaned with Acetone and Iso-propyl alcohol. Step 2: Using shadow masks, 5/100 nm titanium/gold (Ti/Au) is deposited through e-beam evaporation to form a gold coating. The tube is rotated, and Ti/Au is redeposited to ensure metal coating throughout the cylindrical surface of the capillary, leaving a small window at the center along the length of the capillary. Step 3: A miniature thermistor is mounted on the window and soldered to the Au films on either side. Step 4: The capillary tube is anchored onto the IS using silver epoxy. The tube is completely enclosed by the IS at the anchored region using a small copper lid (part of the IS), that has capillary-shaped grooves lined with graphite sheets. Step 5: The capillary and the IS are assembled with the OS using hermetically sealed PEEK tubing and connectors. Step 6: An optical fiber-cum-stopper is inserted into the capillary from one side and is used to localize biological samples near the thermistor while also guiding light for optical imaging. Step 7: The assembled calorimeter is connected to a syringe pump and a physiological buffer is filled in the capillary for biocalorimetry measurements.

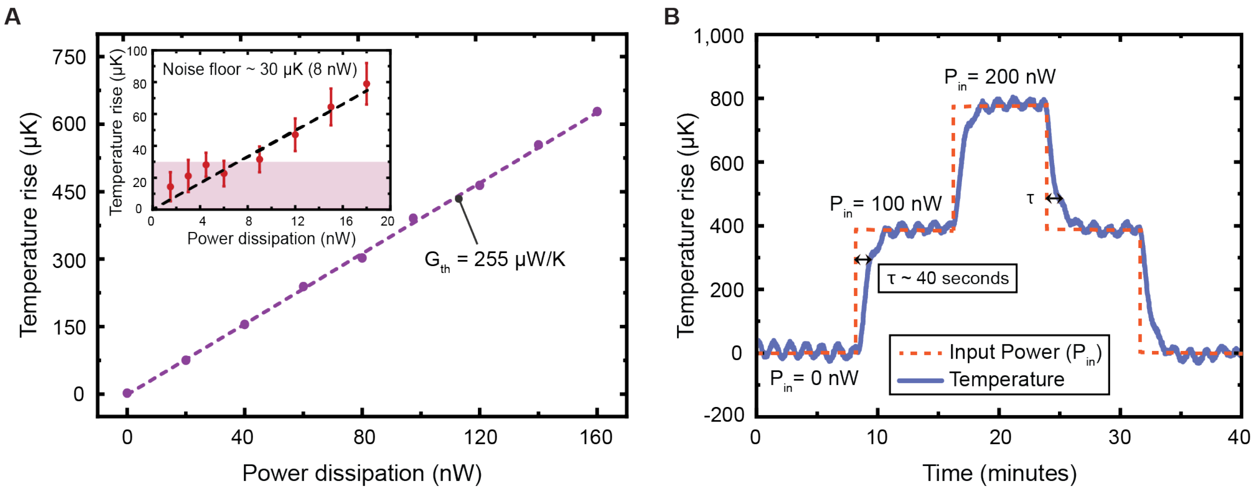


**Fig. S2: Thermal conductance, heat power resolution, and time constant of the calorimeter at 4 µl/min. (A)** The thermal conductance ($G_{th}$) of the sensing capillary to the ambient is ~255 µW/K, in the presence of a continuous buffer flow of 4 µl/min. The temperature measurement noise floor and the corresponding heat power resolution are ~30 µK and ~7.6 nW, respectively (shown in the inset). **(B)** The thermal time constant of the sensing capillary as a response to a constant heat input is ~40 seconds.

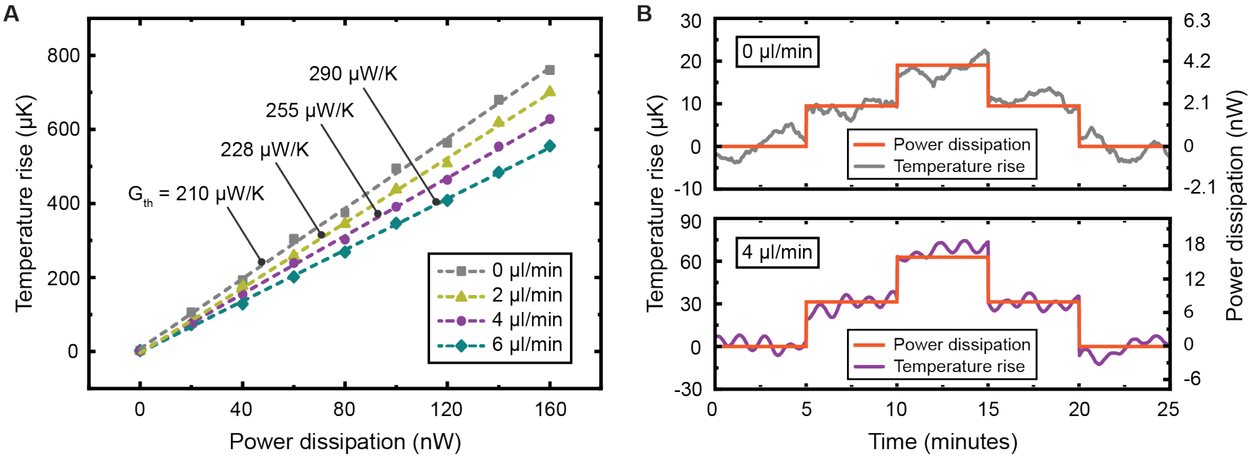


**Fig. S3: Thermal conductance and heat power resolution of the calorimeter at different buffer flow rates. (A)** Measurements of the thermal conductance ($G_{th}$) of the sensing capillary under different flow rates of the buffer medium. $G_{th}$ is 210 µW/K when the tube is filled with buffer but there is no flow. Measured $G_{th}$ are 228 µW/K, 255 µW/K, and 290 µW/K with 2 µl/min, 4 µl/min, and 6 µl/min buffer flow rates, respectively. **(B)** Validation of the thermal resolution of the capillary sensor with no buffer flow and with a 4 µl/min flow rate. The temperature resolution and the heat resolution are ~9.5 µK and ~2 nW, respectively with no buffer flow (i.e. 0 µl/min), and ~30 µK and ~7.6 nW, respectively with a buffer flow rate of 4 µl/min.

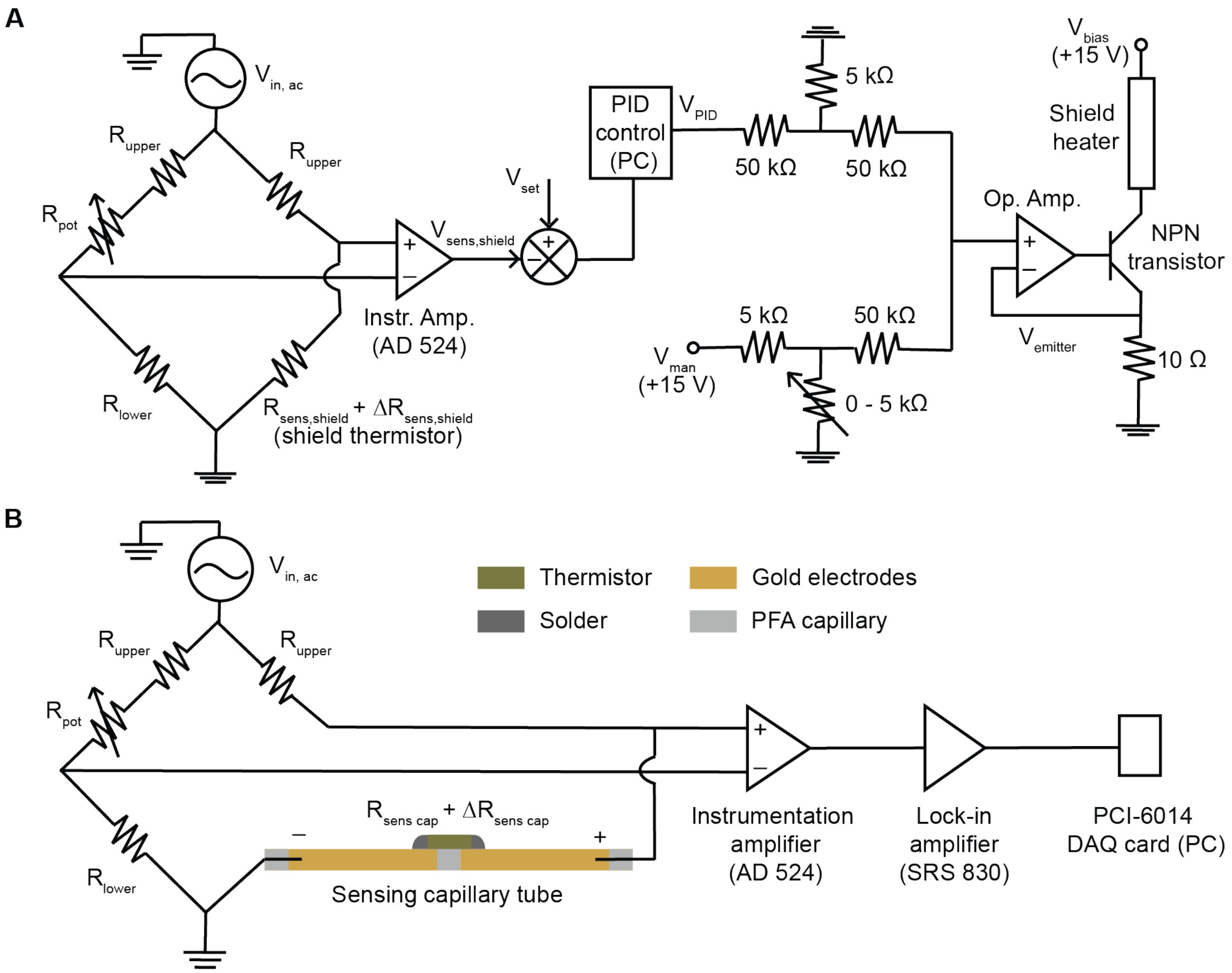


**Fig. S4: Instrumentation for temperature stabilization and for measurement of metabolic heat output. (A)** Circuit diagram for the thermal stabilization of the OS and the IS. AC-driven Wheatstone bridge circuits are used for sensing the real-time temperature fluctuations in the OS and the IS, which is then fed to a PID-based control algorithm that outputs a feedback voltage ($V_{PID}$). This voltage is input to a current-source circuit used to adjust the heating current supplied through the polyimide heaters in the shields to control the Joule heating in the heaters and maintain the shields at the desired temperature set points. **(B)** Circuit diagram for the AC-driven Wheatstone bridge circuit used for high-resolution sensing of the temperature changes due to the metabolic heat output of the biological specimen in the sensing capillary tube. A similar circuit is used for the matching capillary tube to track the common mode temperature drifts.

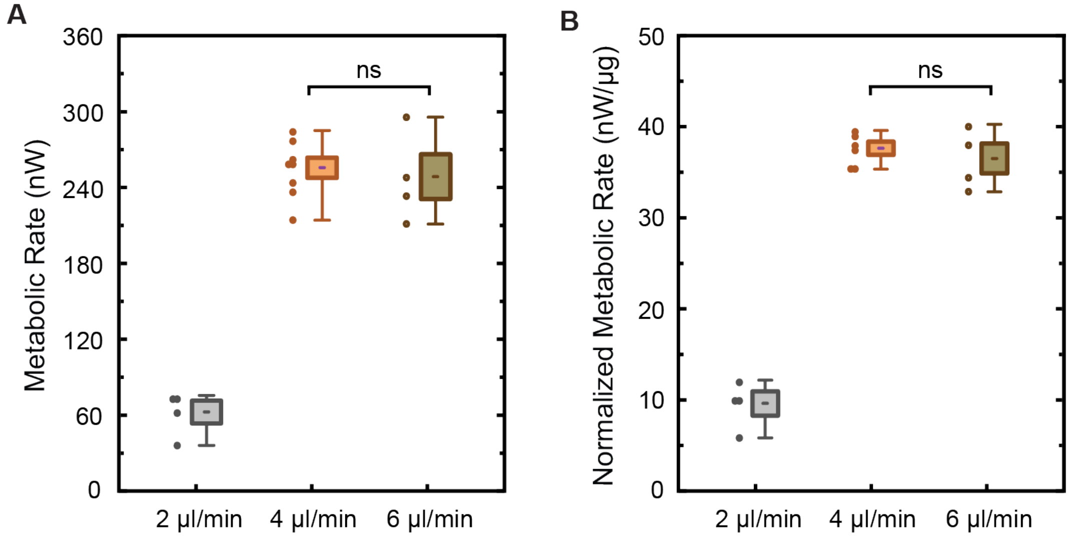


**Fig. S5: Optimization of buffer flow rate for metabolic output measurements. (A)** The distribution of absolute metabolic rates of several brains (10-day-old, female flies, y sc v genotype), measured with buffer flow rates of 2 µl/min, 4 µl/min, and 6 µl/min. **(B)** The distribution of the normalized metabolic rates of the measured brains with the different buffer flow rates. With 4 and 6 µl/min, the observed metabolic rates were stable over 30 minutes showing the brains stay healthy over the period of measurement. With 2 µl/min, the metabolic rates were lower and continuously decreased over a period of 30 minutes, likely due to insufficient supply of oxygen and nutrients to the brains. The distribution plots show each measured data point (solid circles), the mean (dash), the standard error of mean i.e. SEM (shaded box), and the maximum and minimum values (whiskers). A p-value > 0.05 suggests there is no significant (ns) difference between data sets, and $****$, $***$, $**$, and $*$, indicate $p \leq 0.0001$, $p \leq 0.001$, $p \leq 0.01$, and $p \leq 0.05$, respectively.

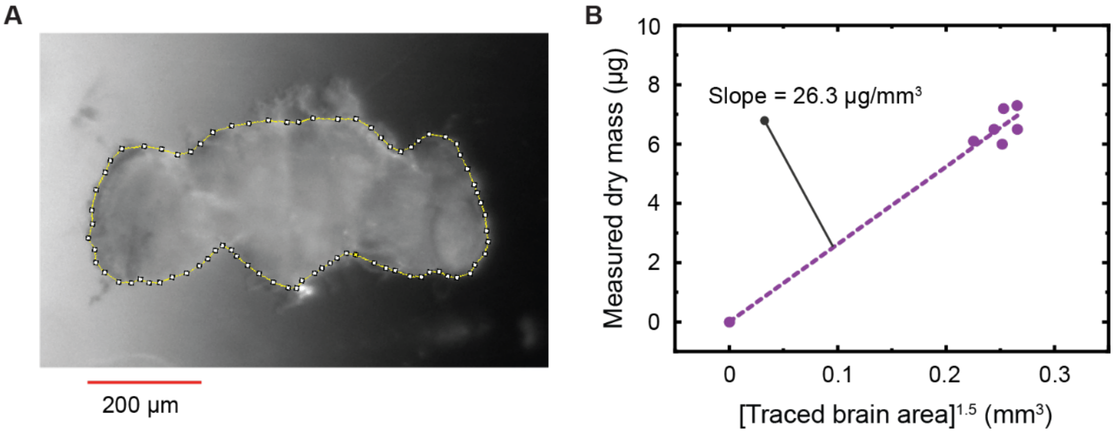


**Fig. S6: Estimation of dry mass of Drosophila brains using optical imaging. (A)** The largest area of cross-section of a brain is traced and measured. **(B)** Measured dry masses for several female brains (10-day-old, y sc v genotype) are plotted against the 1.5$^{th}$ power of their traced area of cross-section, and the proportionality constant is determined to be 26.3 μg/mm$^3$.

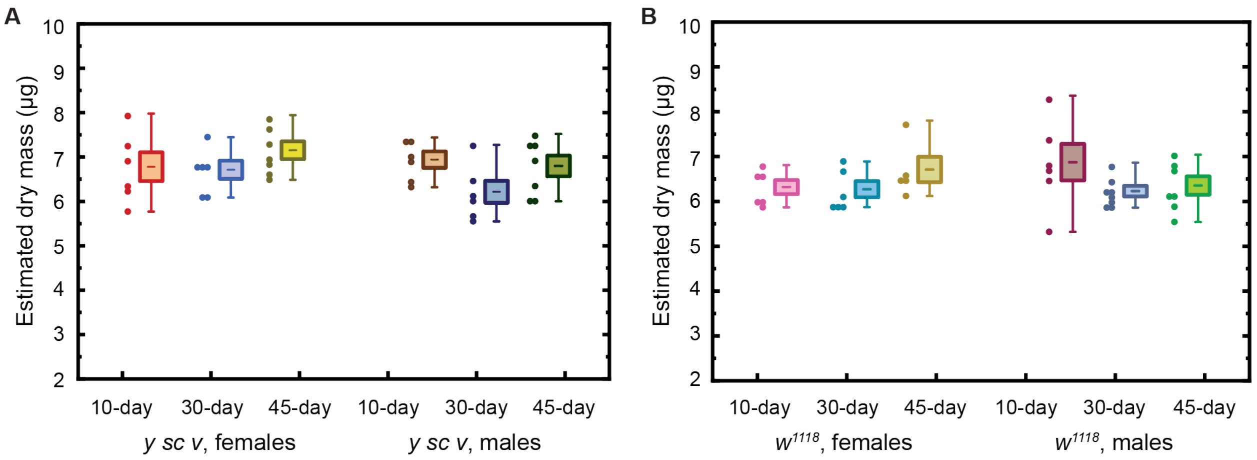


**Fig. S7: Dry mass estimates for all brains used for the gender-, genotype-, and age-related metabolic output measurements. (A)** The estimated dry masses of the male and female brains of the *y sc v* genotype at ages day-10, day-30 and day-45. The metabolic outputs of these brains are reported in Fig. 3C and Fig. 4E. **(B)** The estimated dry masses of the male and female brains of the $w^{1118}$ genotype at ages day-10, day-30 and day-45. The metabolic outputs for these brains are reported in Fig. 3C and Fig. 4F. The above dry masses are estimated using the optical imaging technique shown in fig. S6 and further explained in Methods.

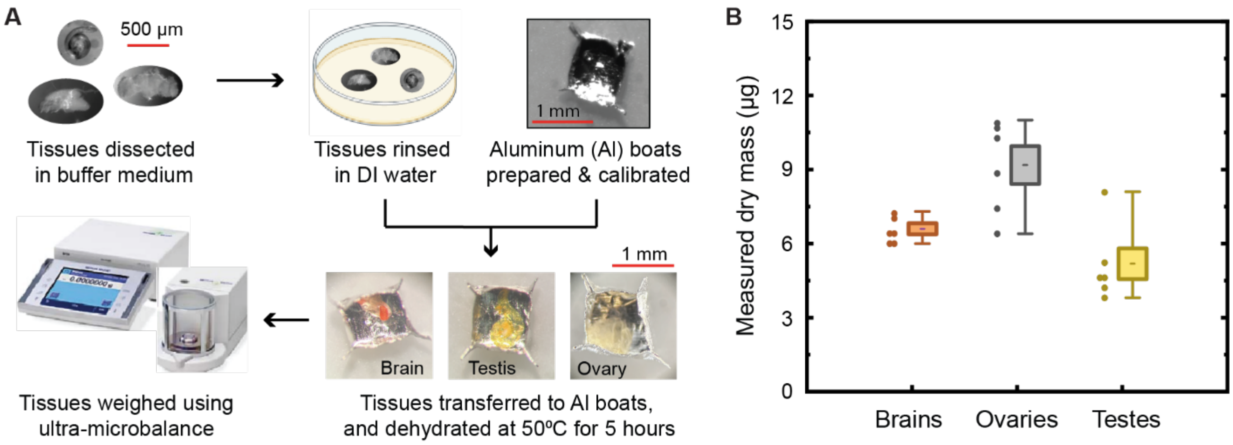


**Fig. S8: Dry mass measurements of Drosophila brains, ovaries, and testes. (A)** Procedure followed to measure the dry mass of Drosophila tissue is illustrated. A tissue is dissected from a fruit fly in the buffer medium, rinsed for 2 – 3 seconds in DI water to get rid of the buffer salts, and then transferred to pre-calibrated weighing boats made of Al foil. The boat with the tissue is then baked at 50ºC for 5 hours, and its dry mass is measured on a Mettler Toledo XP2U ultra-microbalance. **(B)** The distribution of the measured dry masses of several female brains, ovaries, and testes from 10-day-old flies of the y sc v genotype. The mean of the dry masses measured for 6 female brains, 6 ovaries, and 6 testes, are 6.6 µg, 9.2 µg, and 5.2 µg, respectively. The distribution plot shows each measured data point (solid circles), the mean (dash), the standard error of mean i.e. SEM (shaded box), and the maximum and minimum values (whiskers).

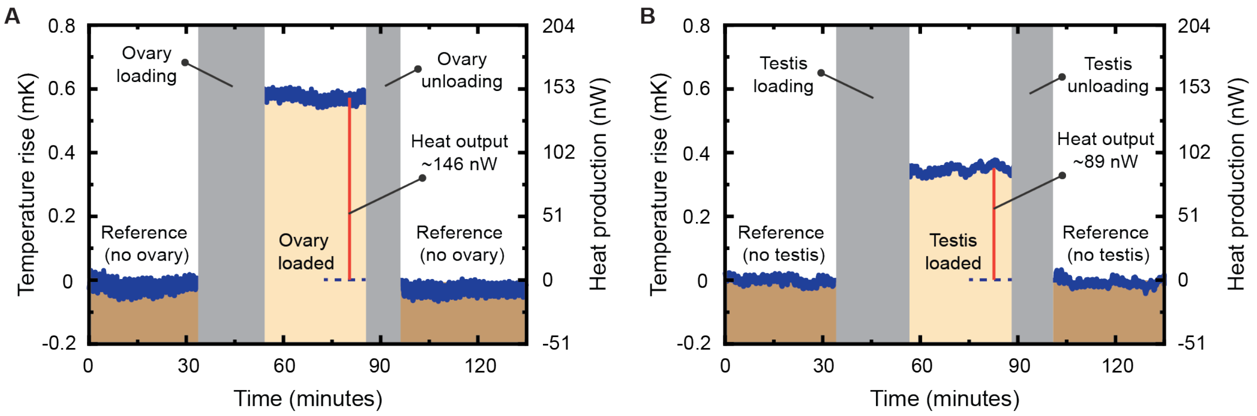


**Fig. S9: Traces of metabolic output measurements of an ovary and a testis. (A)** A trace of the measured metabolic rate from an individual ovary of a female fly (10-day-old, y sc v genotype). **(B)** A trace of the measured metabolic rate from an individual testis of a male fly (10-day-old, y sc v genotype).
